# Seasonal environments drive convergent evolution of a faster pace-of-life in tropical butterflies

**DOI:** 10.1101/2020.05.22.110254

**Authors:** Sridhar Halali, Erik van Bergen, Casper J Breuker, Paul M Brakefield, Oskar Brattström

**Author notes:** **Corresponding author:** Sridhar Halali. **Email addresses:** Erik van Bergen, Casper J Breuker, Paul M Brakefield, Oskar Brattström. **Author contributions:** SH, PMB and OB designed the study; SH carried out the experiment; EvB and OB collected data for habitat preference; CJB wrote a custom ImageJ macro; OB made figure illustrations; SH and EvB carried out analyses and wrote the manuscript with inputs from CJB, PMB and OB. All authors read and approved the final version of the manuscript.

## Abstract

Global change can trigger shifts in habitat stability and shape the evolution of organismal life-history strategies, with unstable habitats typically favouring a faster pace-of-life. We test this hypothesis in species-rich Mycalesina butterflies that have undergone parallel radiations in Africa, Asia, and Madagascar. First, our ancestral state reconstruction of habitat preference, using ~85% of extant species, revealed that early forest-linked lineages began to invade seasonal savannahs during the Late Miocene-Pliocene. Second, rearing replicate pairs of forest and savannah species from the African and Malagasy radiation in a common garden experiment, and utilising published data from the Asian radiation, demonstrated that savannah species consistently develop faster, have smaller bodies, higher fecundity with an earlier investment in reproduction, and reduced longevity, compared to forest species across all three radiations. We argue that time-constraints for reproduction favoured the evolution of a faster pace-of-life in savannah species that facilitated their persistence in seasonal habitats.

## INTRODUCTION

Habitat can act as a templet for the evolution of life-history strategies (Southwood 1977; Southwood 1988). For example, habitats that are heterogeneous in time and/or space and are thus more fluctuating in resource abundance are considered to favour the evolution of ‘fast’ life-history strategies including high growth rates and fecundity with reduced longevity, that allow organisms to rapidly increase population size when opportunities for reproduction are available (Pianka 1970; Southwood 1977). In contrast, selection in more stable habitats is thought to favour ‘slow’ (low growth rates and fecundity with increased longevity) life-history strategies. These fast and slow life-history strategies parallel the dichotomy of *r* and *K*-selected reproductive strategies (Pianka 1970; Gadgil & Solbrig 1972).

The *r/K* selection theory was formulated to predict the type of life-history strategies that would be favoured in colonizing populations with high capacities to increase population size, versus those found in populations that experience density-dependent mortality in stable environments (Pianka 1970; Gadgil & Solbrig 1972). The concept of *r/K* strategies has been criticised for providing a too simplistic overview (see Stearns 1976; Stearns 1992). For example, in many studies, a coarse description of habitat use would suffice as an explanation for the evolution of organismal life-history strategies. In contrast, age-specific demographic models (Gadgil & Bossert 1970; Charlesworth 1980) expand beyond these simple correlations and can therefore be used to help identify causal mechanisms driving the evolution of life-history strategies (Reznick *et al.* 1996; Reznick *et al.* 2002). Nevertheless, comparative analyses using species in close phylogenetic proximity with similar ecological guilds and detailed information on the temporal and spatial characteristics of their habitats can still contribute to understanding the evolution of life-history strategies (Southwood 1988; Partridge & Harvey 1988).

Evolutionary trade-offs are fundamental to optimising investment in life-history traits and can constrain the types of strategies that can be achieved (Stearns 1989; Roff & Fairbairn 2007). One such ubiquitous trade-off is between offspring size and number (Smith & Fretwell 1974). Since internal resources are finite, organisms can either produce fewer offspring of higher quality or many offspring with a relatively low probability of surviving to adulthood (Smith & Fretwell 1974; Van Noordwijk & de Jong 1986). Another well-established trade-off is between voltinism and individual growth rates (Roff 1980; Abrams *et al*. 1996). Insects inhabiting seasonal environments typically only breed during a narrow part of the year when resources for both juveniles and adults are abundant (Tauber *et al.* 1986). The ability to undergo an additional generation in a single breeding season (i.e. multi-voltinism) can dramatically accelerate population growth (Roff 1980; Kivelä *et al.* 2009). To achieve this, there is generally a need to accelerate individual growth rates, which can be achieved by shortening the time needed to complete development, decreasing adult body size, or a combination thereof, resulting in a three-dimensional trade-off between growth rate, development time and body size (Abrams *et al.* 1996; Nylin & Gotthard 1998).

Novel ecological opportunities that may arise under climate change can trigger shifts in habitat preferences and promote adaptive diversification (e.g. MacFadden & Hulbert 1988, Couzens & Prideaux 2018). The origin of tropical savannahs is a dramatic example of a climate-driven biome shift documented in the geological record (Cerling *et al*. 1997; Osborne 2008; Edwards *et al*. 2010). This global expansion of grass-dominated ecosystems during Early-Middle Miocene (11-24 Mya) (Edwards *et al*. 2010) had a major impact on diversification and ecological speciation in both invertebrates and vertebrates, especially in grazing taxa (e.g. MacFadden & Hulbert 1988; Aduse-Poku *et al.* 2009; Couzens & Prideaux 2018). This provides a unique opportunity to study how organismal life-histories have evolved in response to changes in environmental heterogeneity (e.g. Brennan & Keogh 2018). Savannah habitats are highly seasonal with resources for breeding typically limited to a single season (i.e. the wet season), hence imposing strong time-constraints for reproduction. In comparison, tropical forests are relatively stable environments that can sustain resources for breeding, such as grasses for phytophagous insects, throughout the year (Moore 1986; Braby 1995; Halali *et al.* 2020). Given these opposing habitat characteristics, transitions from the ancestral dense canopy forests into newly formed open savannahs are predicted to be particularly challenging for species with short life cycles.

Here we use Mycalesina butterflies to examine how life-history traits have evolved in response to changes in habitat stability. This subtribe of tropical butterflies is distributed across the Old-world tropics and comprises 10 genera (~320-330 species) that have undergone three geographically parallel radiations; one each on the African mainland, Madagascar, and in Asia (Aduse-Poku *et al.* 2015, Figure 1a). Some species of the Asian radiation extend into the sub-tropical regions of north-eastern Australia and others reach as far as Korea and Japan (see Aduse-Poku *et al.* 2015). Focusing on a subset of species from the main African radiation (the genus *Bicyclus),* we recently demonstrated that several independent open-habitat lineages arose during the Late Miocene and Pliocene (Halali *et al*. 2020), and these habitat shifts were likely associated to the expansion of seasonal grasslands during this epoch (Osborne 2008; Edwards *et al*. 2010; van Bergen *et al*. 2016).

**Figure 1:**
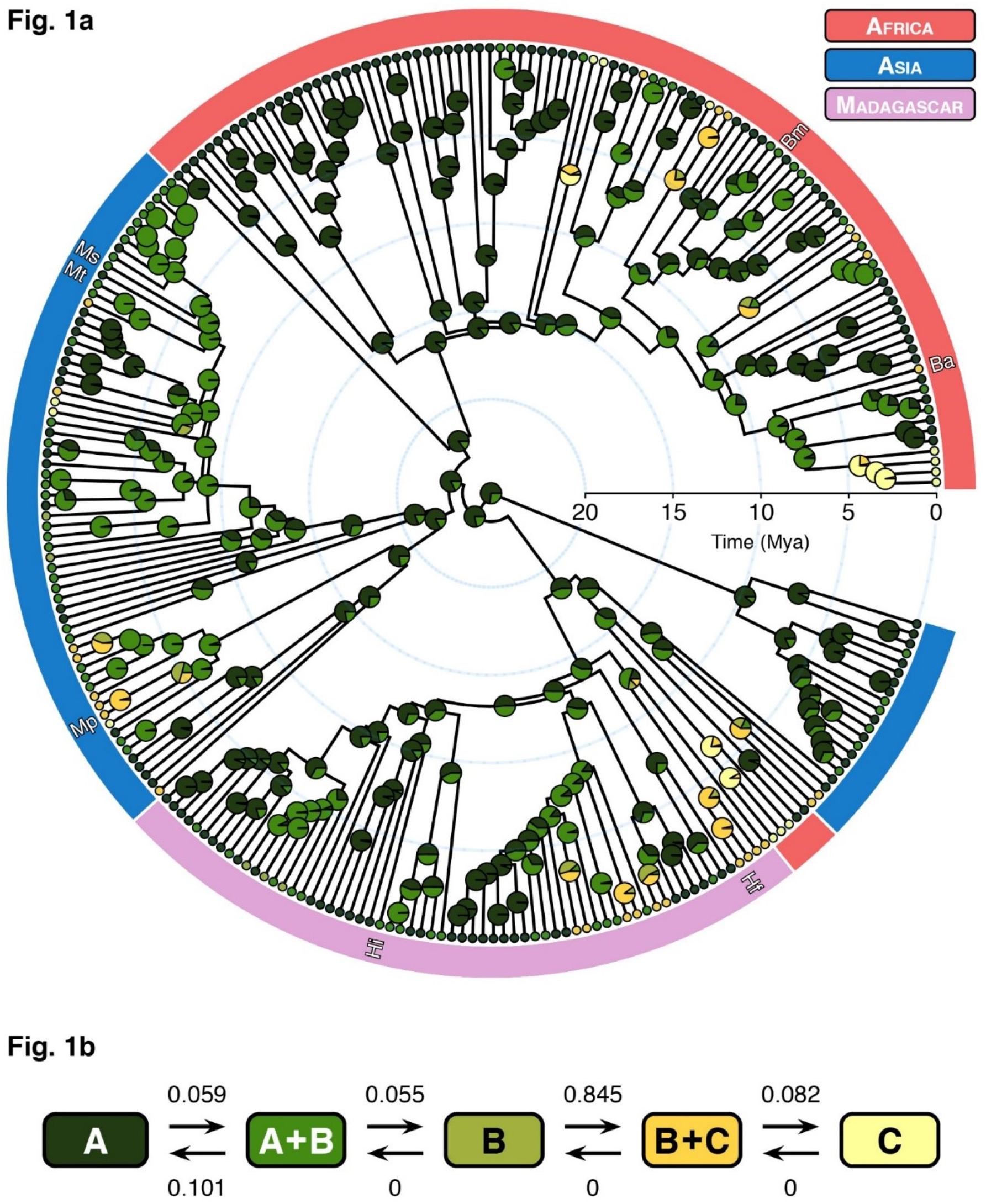
(a) Ancestral state reconstruction for habitat preference in Mycalesina butterflies and (b) the pathway depicting the evolution of habitat specialization with arrows showing transition rates. Letters in the pathway represent habitats categories as follows; A = forests, B = forest-fringes, C = savannahs. Colours in the phylogeny match habitat categories in the evolutionary pathway. Abbreviations marked on the tips of the phylogeny highlight species for which life-history traits were quantified in the common garden experiment (African and Malagasy radiation) and the species from the Asian radiation for which life-history data was available from previously published studies (Ba = *Bicyclus anynana*, Bm = *Bicyclus martius*, Hf = *Heteropsis fraterna*, Hi = *Heteropsis iboina*, Mp = *Mycalesis perseus*, Ms = *Mydosama sirius*, Mt = *Mydosama terminus*).

In this study we broaden the ancestral habitat reconstruction, with near-complete taxa sampling across all geographically independent radiations, to describe patterns of habitat specialization across the entire subtribe of Mycalesina butterflies. We hypothesize that the worldwide expansion of grass-dominated ecosystems also triggered the evolution of open habitat specialists in Asia and on Madagascar, and that transitions occurred in a similar time period. Moreover, from a macroevolutionary perspective, these repeated transitions from stable tropical forests to relatively unstable savannah habitats may have required adaptive changes in a suite of life-history traits. We predict that environmental factors that impede continuous breeding selected for relatively fast life-history strategies in savannah species. To test this hypothesis, we collect life-history data in the laboratory from replicate pairs of forest and savannah species, representing both the African and Malagasy radiation. The availability of data collected previously for three Australasian species (i.e. from the Asian radiation; see Braby & Jones 1994; Braby & Jones 1995; Braby 2002) allowed us to conduct a comparative analysis across all major radiations within Mycalesina butterflies.

Taken together, our results show that Mycalesina butterflies have independently colonised savannahs during the Late Miocene and Pliocene in a stepwise fashion using forest margins and fringes as stepping-stone habitats. Furthermore, the common garden experiments reveal that savannah and forest species have evolved faster and slower life-history strategies, respectively. We argue that the evolution of faster pace-of-life in savannah species is driven by strong time-constraints for breeding in open habitats, and that such life-history strategies contributed to population persistence in seasonal environments.

## MATERIAL AND METHODS

### Ancestral state reconstruction of habitat preference

The evolutionary history of Mycalesina butterflies was reconstructed by classifying the habitat preference for 287 species covering over 85% of all extant taxa and representing all three parallel radiations. Using the available literature, communications with local experts, and our own extensive field experience, species presence was scored in three categories; forest habitats (A), forest-fringes (B), and open or savannah habitats (C). Forest species are restricted to rainforests or habitats with extensive canopy cover. Forest-fringe species are those found in the outskirts of closed-canopy forests or fragmented forests, but never extending into savannahs. Finally, savannah specialists were quantified as species mainly occurring in open grasslands or woodlands where vegetation is dominated by grasses with few interspersed trees. Many species were assigned to two habitat classes (i.e. habitat class A+B or B+C; N=98), and a small number of generalist species were assigned to all three habitats (habitat class A+B+C; N=32). Allowing species to occupy multiple habitats, rather than being fixed to discrete categories, provides a continuum with a decreasing degree of habitat stability; A > A+B > B > B+C > C.

A recent phylogeny from Brattström *et al.* (2020) was used to reconstruct the ancestral states. The evolution of habitat preference was modelled using *fitpolyMk* function in *phytools* ver. 0.7.18 (Revell 2012) which can handle polymorphic states. Since this function can only handle a maximum of two polymorphic states, species that can be found in three habitats (i.e. A+B+C) were dropped, leaving a total of 255 species. We fitted four ordered and four unordered models to the data (i.e. an equal-rates model, a symmetric model, an all-rates-different model, and a transient model, for each model type). The unordered models allow all possible transitions between habitat types, while the ordered models only allow transitions along the habitat continuum. Model performances were assessed using the Akaike Information Criteria (AIC) score and an ancestral state reconstruction was conducted for the best fitting model using Bayesian stochastic mapping (Bollback 2006) implemented in *phytools* using *make.simmap* function. We simulated 1000 character states on the Maximum Clade Credibility tree and summarised the states at each node to indicate the probability of a particular state to be ancestral.

### Common garden rearing

Laboratory populations of replicate pairs of forest and savannah species from the African (*Bicyclus spp*.) and the Malagasy (*Heteropsis spp*.) radiations were used to explore the role of seasonality in shaping the evolution of life-history strategies. Species inhabiting the stable African forests were represented by a laboratory population of *Bicyclus martius* that was established in 2018 from gravid females collected in Bonkro, Ghana. This species is strongly linked to intact rainforests in West Africa but can also be found in fairly open clearings and forest margins as long as some canopy cover remains (habitat class A+B; see Larsen 2005; Oostra *et al*. 2014a). Species inhabiting the more seasonal habitats on the African mainland were represented by a population of *Bicyclus anynana* from Nkhata Bay in Malawi (Brakefield *et al*. 2009). This extensively studied species is widely distributed across open woodland and savannah habitats in East Africa and never extends into forests with complete canopy cover (habitat class B+C; Larsen 1991; Windig *et al*. 1994). The Malagasy radiation was represented by *Heteropsis iboina* (forest; class A+B) and *H. fraterna* (savannah; class B+C). The laboratory stock of *H. iboina* was established in 2013 from gravid females collected in Andasibe-Mantadia NP in Madagascar (van Bergen *et al*. 2017), while *H. fraterna* was established in 2018 from females collected in Ranomafana NP (Madagascar). All laboratory stocks were maintained at 25°C and 70% relative humidity (RH) under a 12:12 hr L:D photoperiod resembling wet season conditions in the field. Young maize plants (*Zea mays*) were host plants for *B. anynana,* wheat plants (*Triticum aestivum*) for *H. fraterna,* and running mountain grass (*Oplismenus compositus*) for *B. martius* and *H. iboina. Oplismenus spp*. are natural hosts for many Mycalesina butterflies, and all stock populations demonstrate high performance on *Oplismenus* grasses (e.g. Kooi 1992; Oostra *et al*. 2014; Nokelainen *et al*. 2016).

### Measuring life-history traits

Eggs were collected from each laboratory population at 24-hour intervals (between 9.30 to 10.30 hours), from small *Oplismenus* plants and then transferred to petri dishes lined with a filter paper and kept in climate-controlled chambers (Sanyo/Panasonic MLR-350H) at 25°C and 70% RH under a 12:12 hour L:D photoperiod. The eggs of each species were photographed using a Leica DFC495 camera coupled to a Leica M125 stereoscope, and the cross-sectional area of each egg measured from the resulting images in ImageJ (Schneider *et al*. 2012) using a customised macro. Areas of eggs measured in this way are highly correlated with egg weight (e.g. Gibbs *et al*. 2010). The egg development time was estimated as the number of days between laying date and hatching date.

Upon hatching, three cohorts of 60 newly hatched larvae (N=180) for each species were randomly allocated to large rearing cages (35 × 46 × 60 cm). Larval host plants (*Oplismenus* grass) were watered daily and replaced when necessary to ensure ad libitum feeding and the position of the cages within the climate-controlled chambers was changed daily. Larval development time was the number of days from egg hatching to pupation; pupal development time, the number of days from pupation until adult eclosion; and total development time, the sum of larval and pupal development times. All pupae were weighed to the nearest 0.1 mg (Mettler AE163) within 24 hours of pupation. Larval growth rates were calculated by dividing the natural log of the pupal weight by the larval development time (see Gotthard *et al*. 1994). After eclosion, two to four day old virgin females (except two females which were one day old) were allowed to mate with virgin males to determine daily variation in fecundity and to compute individual fecundity curves. For *B. anynana, H. iboina,* and *H. fraterna,* we were able to select 15 mating pairs in the first trial. For *B. martius,* we could not obtain any mating pairs in the first trial, and we only observed a single mating after multiple trials. Therefore, males and females of this species were kept in a single cage for approximately 15 days, after which females were randomly selected for measuring fecundity. After copulation, individual females were kept in cylindrical plastic pots (11.5 × 13.5 cm) and provided with a cutting of *Oplismenus* grass kept in water for oviposition. The number of eggs laid by the females was assessed every 24-hours for 15 consecutive days. The females of *B. martius* were dissected after the monitoring period to confirm their mating status. Three non-mated individuals were excluded from the data set yielding a total sample size of four for this species. Female longevity was measured as the percentage of females that were alive after the 15-day fecundity assessment period. For *B. martius* we included both non-mated and mated females (N=7). All females were fed on slices of moist banana throughout the experiment.

### Statistical analyses

Generalised linear models (GLM) were used to examine the effects of habitat class, genus, sex, and their interactions, on a suite of life-history traits. Habitat class was a categorical variable (forest or savannah), and genus (*Bicyclus* or *Heteropsis*) was included to account for phylogenetic relatedness between the species. Sex was excluded from the models for egg size and egg development time since this variable cannot be established at this life stage. Data on development times (egg, larval, pupal and total) were analysed using GLMs with a Poisson distribution and a log link function. GLMs with a Gaussian distribution were used to fit the data for egg size, pupal weight and growth rate. Post hoc pairwise comparisons (Tukey’s HSD; α = 0.05) were carried out using *emmeans* R package (Lenth 2019).

Individual fecundity curves were estimated using a generalised linear mixed model with a negative binomial distribution, as implemented in the R package *glmmTMB* (Magnusson *et al*. 2019). In addition to habitat class and genus, the number of days (centred) and all interactive terms were included as fixed effects. Pupal weight, centred within each species, was added as a covariate. Finally, since the raw data suggested that the fecundity curves had non-linear distribution, we included a quadratic term for the number of days in the model. We initially fitted full models, allowing interactions between all predictors, and the minimum adequate model was found by AIC guided backward elimination. All statistical analyses were performed in R ver. 3.6.1 “Action of the Toes” (R core team 2019).

## RESULTS

### Ancestral state reconstruction for habitat preference

An ordered all-rates-different model provided the best fit for the habitat preference data (see Table S1 in Supporting Information). This model suggested that the lineages that gave rise to three geographically independent radiations were forest specialists and that more open habitats were colonised repeatedly during the Late Miocene and Pliocene (8-3 Mya; Figure 1a). The transition matrix of the best-fit-model suggests that forest fringe habitats represent a stepping-stone towards adaptation to strictly open environments (Figure 1b). In other words, the invasion of woodland and savannah habitats were preceded by adaptation to semi-shaded habitats. Moreover, parameter estimates suggest that back-transitions, that is transitions from open to forest, are unlikely (Figure 1b).

### Egg size and development

We found that forest species laid larger eggs and that their offspring took longer to complete embryogenesis (habitat, P<0.001 for both egg size and egg development time; Figure 2). The effect of habitat-use was larger in the species representing the African radiation than in those representing the Malagasy radiation (habitat:genus, P<0.05 for both traits; see Table S2 & S4). The results from our study are in line with the habitat-dependent patterns observed for the Australasian species from the Asian radiation (Braby & Jones 1994; Braby & Jones 1995). Here, compared to savannah species (*Mycalesis persons*), the two forest specialists (*Mydosoma sirius* and *Mydosama terminus*) laid larger eggs that took longer to develop (data from Braby & Jones (1994) are presented in Figure 2; see Supporting Information for details). Note that all three species were placed in the genus *Mycalesis* at the time of the original publication.

**Figure 2:**
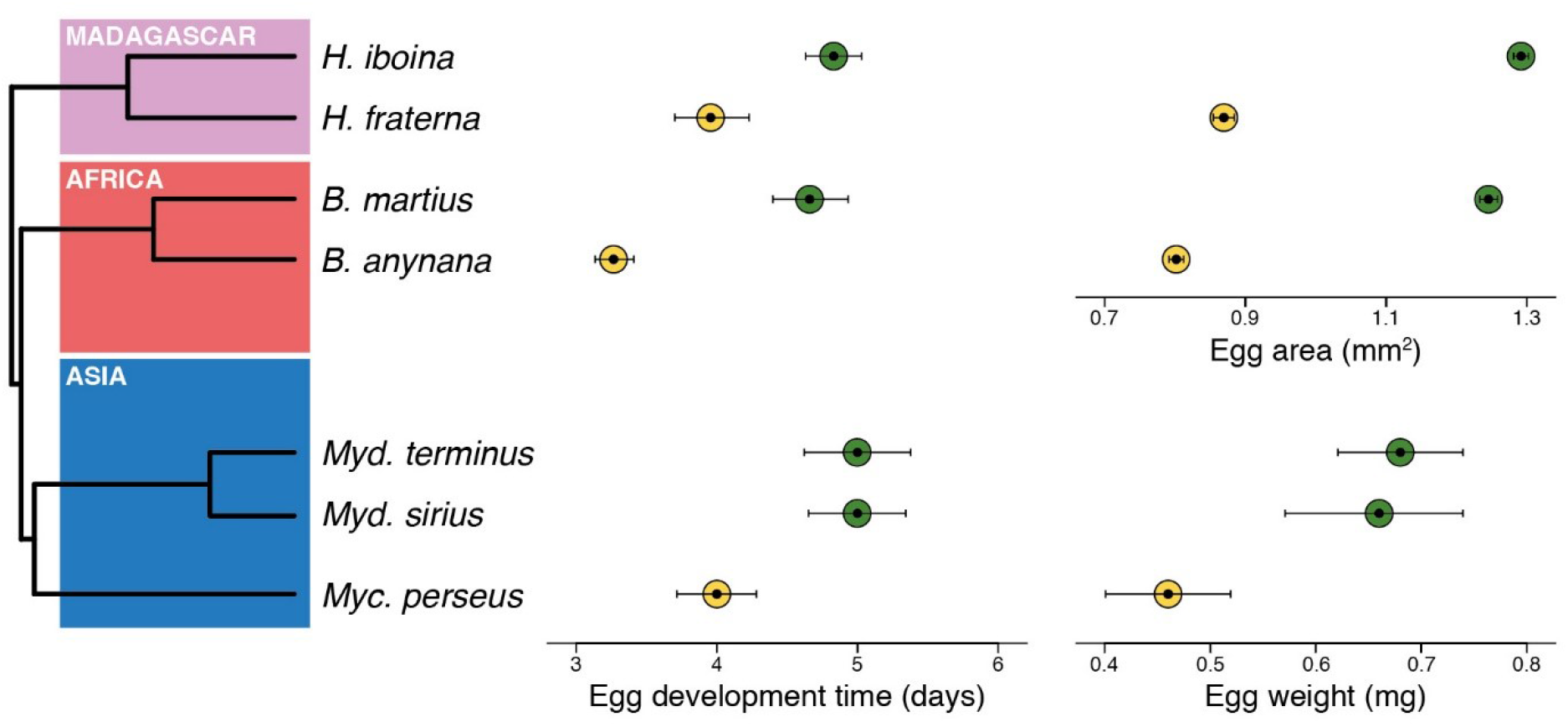
Phylogenetic relationships among the species included in the study which represent all three radiations, together with mean values and 95% confidence intervals for egg development time and egg size. Forest and savannah species are shown in green and yellow circles, respectively. For the African and Malagasy radiation, savannah species had significantly reduced egg sizes and egg development times (see Results). The data on species from the Asian radiation were extracted from Braby & Jones (1994) (see Supporting Information) and error bars for the egg weights represent standard deviations.

### Development times and growth rates

Compared to savannah species, forest species had longer larval, pupal and total development times (habitat, P<0.001 for all traits; Figure 3a; figures for larval and pupal development time in Figure S1a & S1b). Differences in developmental times between habitat specialists were larger in the species representing the African radiation (habitat:genus, P<0.002 for all traits; see Table S5-S7). Forest species had lower individual growth rates (habitat, P<0.001) and a larger body mass (habitat, P<0.001) than savannah species (Figure 3b & 3c; Table S8 & S9). All species were sexually dimorphic for body size (all pairwise comparisons, P<0.001) with female pupae weighing on average 22% more than males, except for *B. martius* where males were slightly heavier than females. These results complement the data on the Australian Mycalesina butterflies from the Asian radiation where two forest species had longer total development times, lower growth rates and larger body sizes than the savannah specialist (data from Braby & Jones (1994) are presented in Figure S3).

**Figure 3:**
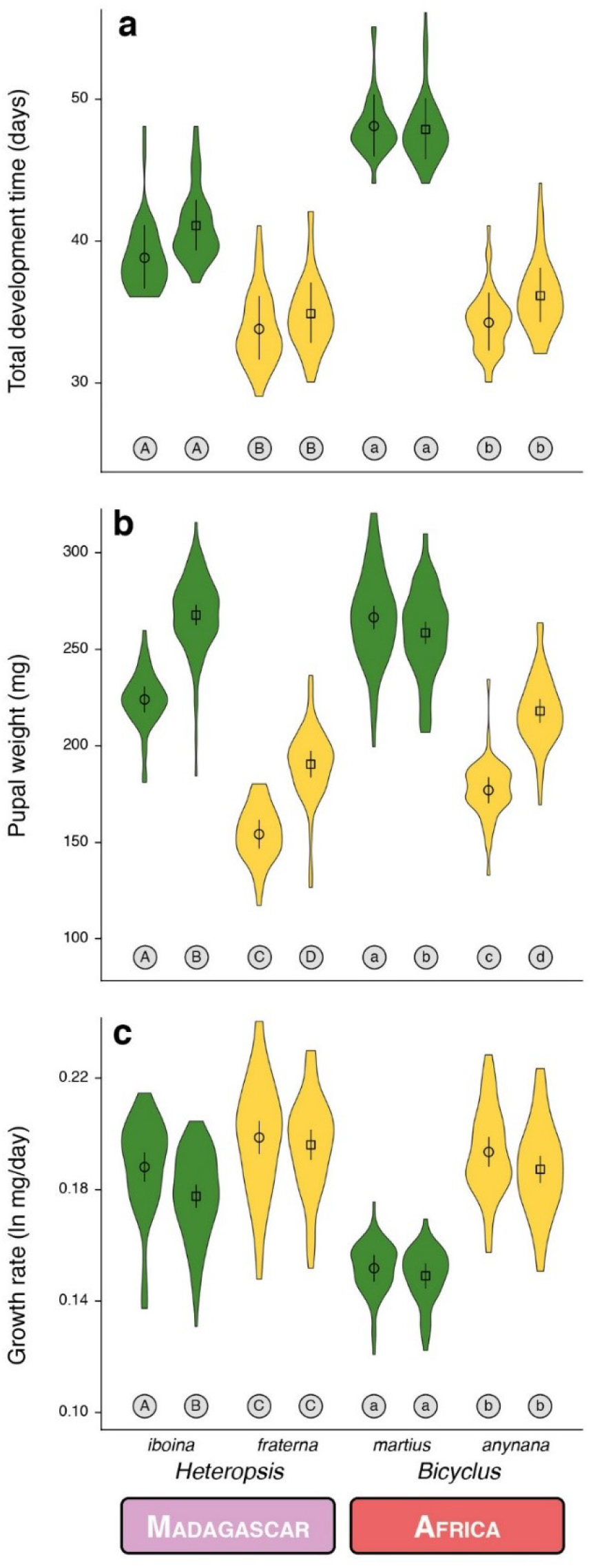
Violin plots with mean and 95% confidence intervals for the (a) total development time, (b) pupal weight, and (c) growth rate for replicate pairs of forest and savannah species from the Malagasy and African radiation reared in the common garden experiment (green = forest species, yellow = savannah species). Sexes are denoted with different shapes (circles = males, squares = females) and significant differences between groups (Tukey’s HSD, P < 0.05) are indicated by different letters, coding for each radiation independently. Data for these life-history traits for species from the Asian radiation are presented in Figure S2.

### Fecundity curves and longevity

In both the African and Malagasy radiations, the forest species had lower fecundity and laid eggs more uniformly across their lifespan than the savannah species (habitat, P<0.001; Figure 4). The magnitude of these differences appeared to be slightly higher in the African than in the Malagasy radiation. For example, the African savannah species (*B. anynana*) had higher fecundity and a more pronounced early investment than the savannah species from the Malagasy radiation (*H. fraterna*) (Figure 4). In contrast, the fecundity curves of both forest species were closely similar. After 15 days of fecundity measurements, 55% females of forest species and 6% of savannah species survived in the African radiation (Figure 4). Similarly, 46% and 20% of forest and savannah species survived in the Malagasy radiation, respectively (Figure 4). Note that the females from the forest species from Africa, *B. martius*, were kept in communal cages for 15 days prior to fecundity assessments (see Methods). The high proportion of females still alive after the assessment suggests that this species is potentially extremely long-lived. Similarly, in the Australian species, the savannah species had higher fecundity and a prominent early peak in fecundity compared to two forest species (data from Braby & Jones (1995) are presented in Figure S4). Systematic data on longevity were not available for these species.

**Figure 4:**
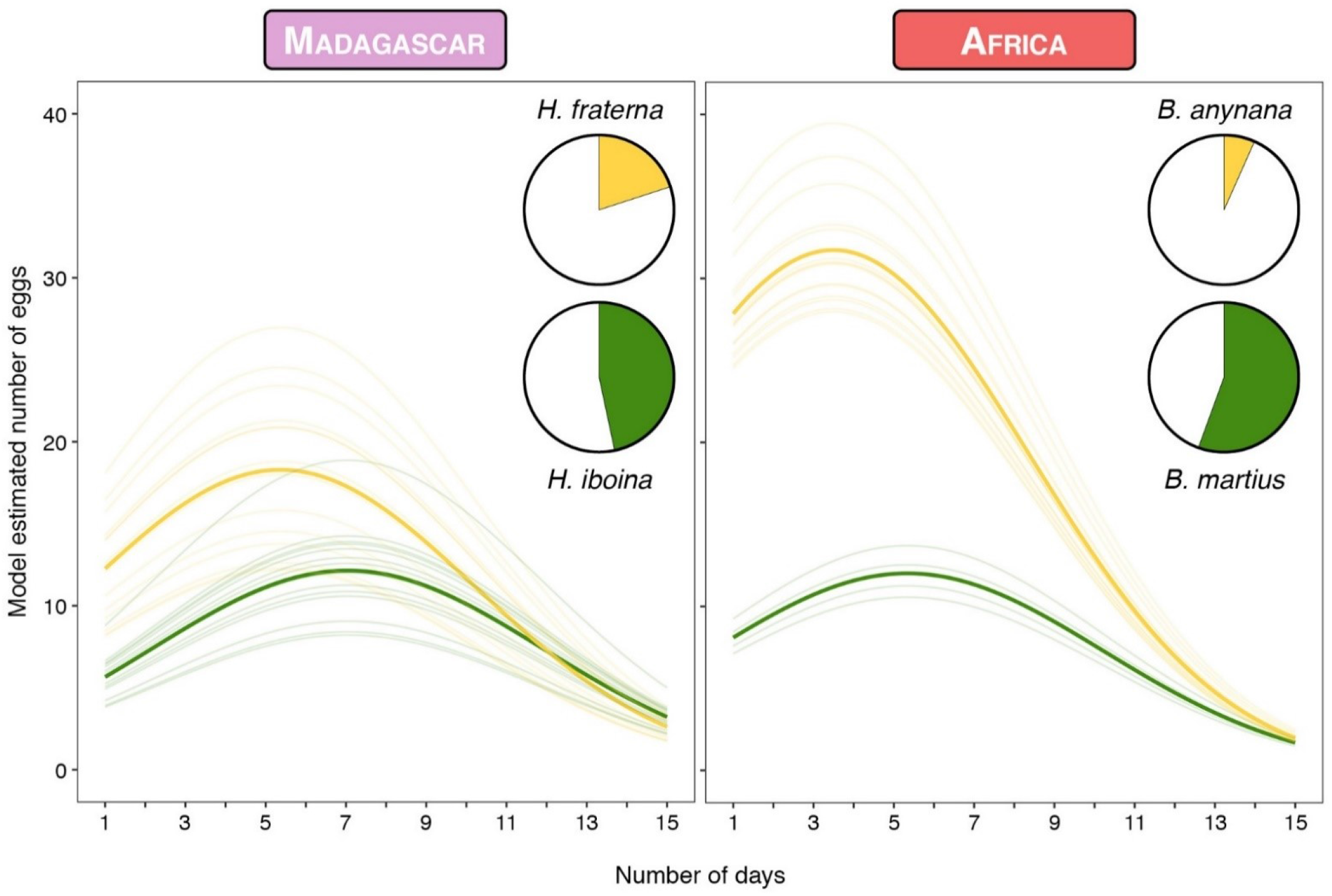
Fecundity curves for replicate pairs of forest and savannah species from the Malagasy and African radiation (green = forest species, yellow = savannah species). Thick and thin lines represent the model estimated average and individual fecundity curves, respectively. In the inset, highlighted portions of pie charts show the percentage of surviving females after 15 days of fecundity assessment. Fecundity curves for species from the Asian radiation are presented in Figure S4.

## DISCUSSION

Adaptive radiations comprise the rapid differentiation of a single common ancestor into an array of species that inhabit a variety of environments and differ in the phenotypic traits used to exploit these ecological niches (Losos & Mahler 2010; Stroud & Losos 2016; Gillespie *et al*. 2020). The process includes both ecological diversification and speciation (Schluter 2000; Nosil 2012), and a striking feature of this phenomenon is that parallel radiations frequently produce convergent forms. For example, *Anolis* lizards have evolved closely similar series of ecomorphs and ecomorph-specific morphologies on numerous Caribbean islands (Losos *et al*. 1998; Mahler *et al*. 2013), benthic and limnetic ecotypes of stickleback fish have emerged repeatedly in postglacial lakes (Schluter & Nagel 1995; Rundle *et al*. 2000), and similar habitat-specific morphological features have evolved multiple times in cichlid fishes, both within and across lakes (Kocher *et al*. 1993, Muschick *et al*. 2012). Our results demonstrate that colonisation of open seasonal habitats occurred repeatedly in the three geographically parallel radiations of Mycalesina butterflies and that these transitions are associated with the convergent evolution of life-history strategies.

The global expansion of open woodland and savannah habitats during the Miocene is an important driver of ecological diversification in a variety of herbivorous taxa including both invertebrates (e.g. Zarlenga *et al*. 2006; Aduse-Poku *et al*. 2009; Kergoat *et al*. 2018) and vertebrates (e.g. MacFadden & Hulbert 1988; Couzens & Prideaux 2018; Fuchs *et al*. 2019). Our results suggest that this climate-driven biome shift also played a crucial role in the evolutionary history of Mycalesina butterflies (see also van Bergen *et al*. 2016; Halali *et al*. 2020). During this epoch, much of the trans-continental dense canopy forest was replaced by a mosaic of open woodland and savannah habitats (Jacobs *et al*. 1999; Jacobs 2004; Edwards *et al*. 2010), and this heterogeneous environment appears to have unlocked ecological opportunities for Mycalesina butterflies. Our ancestral state reconstruction for habitat preference suggests that colonisation of open habitats was preceded by adaptations to semi-shaded habitats, such as forest clearings or margins. Such a ‘stepping-stone’ model of habitat preference evolution has been described for a variety of other taxa (e.g. Ruck *et al*. 2006, see examples in Donoghue & Edwards 2014), implying that direct transitions to more extreme habitats may be rare, and often require pre-adaptations or evolution of key traits which facilitate persistence in harsh environments (Donoghue & Edwards 2014).

Tropical forests and savannahs differ considerably in their degree of seasonality. Closed-canopy forests can buffer extreme seasonal differences (for example in temperature or humidity; Montejo-Kovacevich *et al*. 2020) and can sustain vital resources, including grasses for grass-feeding insects, throughout the year (Moore 1986; Braby 1995). In contrast, tropical grasslands are highly seasonal environments where resources for reproduction are only present during a short period of the year. The convergent evolution of life-history traits within this adaptive radiation of tropical insects is likely to be associated with these habitat-specific differences in seasonality. The habitat templet model (Southwood 1977; Southwood 1988) postulates that selection will favour slow strategies in species inhabiting stable habitats with continuous or prolonged breeding seasons, while fast strategies will occur in habitats with strong fluctuations in resource abundance where animals experience strong time-constraints for breeding. Long-term data from Malawi has revealed that savannah species only reproduce during the wet season (~five months) and may approximately fit two or more generations before undergoing reproductive diapause to cope with the lack of larval host plants in the dry season (Halali *et al*. 2020). In contrast, forest species reproduce throughout the year, suggesting that time-constraints for reproduction are stronger for savannah species (Halali *et al*. 2020). We have shown that savannah species of Mycalesina butterflies have shorter development times with higher growth rates and hypothesize that this strategy allows these species to increase the number of generations they can fit into a single breeding season even if it results in lower critical body mass.

Time-constraints on reproduction may also have shaped patterns of reproductive effort across species. Theoretical studies predict that short-lived individuals with high early fecundity are favoured by selection in time-constrained environments compared to habitats where such constraints are absent or weak (Southwood 1977; Kivelä *et al*. 2009). Our fecundity estimates reveal that savannah species have higher reproductive effort early in their lifespan, while the distribution is more uniform in forest species. Furthermore, there is a trade-off between early reproductive effort and longevity (Miyatake 1997; Zera & Harshman 2001; Jervis *et al*. 2007), with larger bodied insects typically living longer and having higher fecundity (Honěk 1993; Nylin & Gotthard 1998; Holm *et al*. 2016). Indeed, we find that forest species are generally longer lived with larger body sizes than Mycalesina species from savannah habitats. Also, another forest species *Bicyclus sanaos*, a sister species of *B. martius*, has a significantly longer lifespan compared to the savannah species *B. anynana* (Oostra *et al*. 2014a).

Another ubiquitous life-history trade-off is between egg size and number (Smith & Fretwell 1974). Since internal reserves for reproduction are limited (van Noordwijk & de Jong 1986), increased fecundity is typically associated with a reduction in egg size, and vice versa (Smith & Fretwell 1974; Berrigan 1991). Our data reveal that forest species produce relatively few eggs of high quality (i.e. greater egg area), while closely related savannah specialists produce more eggs that are much smaller in size. Similar habitat-dependent trade-offs between egg size and number were demonstrated in a comparative study of temperate forest and open habitat Satyrine butterfly species (Karlsson & Wiklund 2005). Since egg size is generally positively correlated with female body size in insects (Berrigan 1991, García-Barros 2000) and forest species usually have larger body sizes than savannah species (e.g. Braby 2002), smaller eggs in the latter could also be a consequence of a reduction in body size (e.g. Davis *et al*. 2011). In our study, the differences in egg size and fecundity among habitat specialists remained significant after accounting for variation in body size (Table S3 & Figure 4). As a potential caveat we note that the fecundity of the African forest species *B. martius* was measured 15 days post-eclosion, and the females may have laid some eggs before the fecundity assessment. However, since this species is extremely long-lived (see Results and Oostra *et al*. 2014a) we are confident that our fecundity measurement only represents a slight underestimate.

Apart from trade-offs, including those resulting from time-constraints, hypotheses related to adaptive foraging could potentially also explain the observed differences in egg size across species. The size of the eggs is strongly correlated to the size of the larval head capsules, which has been shown to affect the foraging ability of early instar larvae (Braby 1994, Massey & Hartley 2009). Braby (1994) postulated that butterflies using less palatable host plants may lay larger eggs hence produce hatchling with larger head capsules. This adaptation could maximise the survival rate of early instar larvae on hosts with high physical resistance and/or those with low nutritional values. Empirical evidence supporting this hypothesis was provided by experimental work using Australian Mycalesina (Braby 1994) and Japanese Hesperiidae (Nakasuji 1987) butterflies. The habitat-dependent differences in egg size observed in our experiment are unlikely to be in line with the predictions of the adaptive foraging hypothesis. Today’s tropical savannahs and grasslands are dominated with grasses using the C4 photosynthetic pathway (Osborne & Beerling 2006; Edwards *et al*. 2010) that typically have high fibre levels (i.e. greater toughness) and low levels of nutritional quality (Barbehenn *et al*. 2004a; 2004b). Since open-habitat species are more likely to use these host plants of lower palatability (van Bergen *et al*. 2016; Nokelainen *et al*. 2016), selection on the foraging ability of early instar larvae would have favoured the evolution of larger egg sizes in open-habitat Mycalesina species. Instead, our data revealed that savannah species produce relatively small eggs.

Ectotherms inhabiting unstable or seasonal environments often demonstrate developmental plasticity to cope with environmental heterogeneity (e.g. Beldade *et al*. 2011; Kivelä *et al*. 2013). This environmental regulation of development can result in a better match between adult phenotype and adult environment and typically involves the concerted response of a suite of traits to external conditions (Parsons & Robinson 2006; van Bergen *et al*. 2017). Mycalesina butterflies, particularly savannah species, demonstrate plasticity in morphological (e.g. Windig *et al*. 1994), physiological (e.g. Oostra *et al*. 2011), behavioural (e.g. van Bergen & Beldade 2019) and life history traits (e.g. Oostra *et al*. 2014b). Individuals that fly during the wet season reproduce actively, have smaller body size and shorter lifespan, while those emerging in the dry season undergo reproductive diapause, have larger body size and longer lifespan (Brakefield & Reitsma 1991, Halali *et al*. 2020). In this study we focussed on the life-history traits of reproductively active adults by rearing cohorts at temperatures that mirror the conditions of the wet season. The evolution of plasticity itself may also have played a crucial role in the colonisation of seasonal environments by Mycalesina butterflies and is the topic of future work.

Finally, variation in extrinsic mortality risk may play a role in the evolution of life-history traits across habitats. In seasonal environments, the availability of resources that are utilized by prey species is typically highly synchronised with the abundance of parasitoids and predator communities (Morais *et al*. 1999, Molleman *et al*. 2016). The wet season in open habitats may therefore represent a high predation risk environment where the production of many small eggs and investment in early reproduction is predicted to be advantageous (e.g., Reznick & Endler 1982; Reznick *et al*. 1990). Moreover, faster larval growth rates may require longer or more frequent foraging bouts, which could increase the predation risk of savannah species (Johansson & Rowe 1999; Gotthard 2000). In contrast, the prolonged development of forest species could potentially increase their exposure to natural enemies (Benrey & Denno 1997; Williams 1999). Overall, risks of enemy attacks in insects have been shown to be lower in closed-canopy forests compared to semi-open habitats such as forest clearings (Richards & Coley 2007). Future studies focussing on quantifying age-specific mortality (e.g. Reznick *et al*. 1996) are needed to improve our understanding on the role of extrinsic mortality on the evolution of life-history strategies in Mycalesina butterflies.

The parallel nature of three independent and species-rich radiations of Mycalesina butterflies allowed us to extend our results beyond simple correlation between environment stability and associated life-history strategies. We show that even with considerable periods of independent evolution, *r*-selected Mycalesina butterflies have evolved repeatedly in seasonal savannahs, suggesting that the faster pace-of-life was required to be able to persist in these unstable habitats (see Pianka 1970; Southwood 1977; Braby 2002). As recently found in parallel for the evolution of reproductive diapause (Halali *et al*. 2020), we argue that time-constraints on reproduction act as a strong selective agent in these open habitats and that trade-offs among life-history traits played a central role in the evolution of life-history strategies in these tropical butterflies.

## ACKNOWLEDGEMENTS

We would like to thank Chris Müller, David Lees, David Lohman, John Tennent, Peter Roos, Richard Vain-Wright, Sáfián Szabolcs, Steve C. Collins, Ullasa Kodandaramaiah, and Wenda Cheng, for helping with habitat classification; Jack Thorley for statistical advice and Melanie Gibbs for fruitful discussions on butterfly reproductive strategies. Financial support was provided by grants from the European Research Council (EMARES, Grant No. 250325) and the John Templeton Foundation (Grant No. 60501) to PMB. SH was supported by the John Stanley Gardiner Studentship.

## SUPPLEMENTARY INFORMATION

**Table S1:** AIC scores of the four ordered and four unordered models fitted using *fitpolyMk* function in *phytools*. The best fitting model is highlighted in bold and was used for reconstructing the ancestral states presented in Figure 1 of the main text.

| Model | Ordered | AIC |
| --- | --- | --- |
| Equal rates | No | 617.11 |
| Equal rates | Yes | 533.84 |
| Symmetric | No | 521.14 |
| Symmetric | Yes | 510.47 |
| All rates different | No | 498.47 |
| **All rates different** | **Yes** | **480.23** |
| Transient | No | 596.06 |
| Transient | Yes | 535.84 |

**Table S2:**
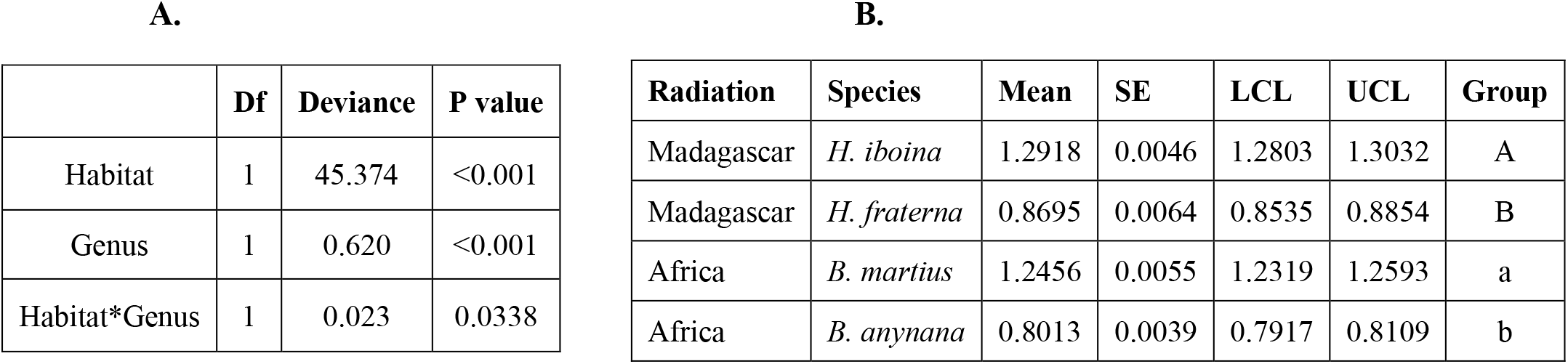
(A) Analysis of deviance table for egg area and (B) model estimated marginal means with upper and lower confidence limits. Significant differences (Tukey’s HSD, α = 0.05) between species were obtained using the R package *emmeans* and are indicated by different letters coding for each radiation independently.

|  | Df | Deviance | P value |
| --- | --- | --- | --- |
| Habitat | 1 | 45.374 | <0.001 |
| Genus | 1 | 0.620 | <0.001 |
| Habitat*Genus | 1 | 0.023 | 0.0338 |

B.
| Radiation | Species | Mean | SE | LCL | UCL | Group |
| --- | --- | --- | --- | --- | --- | --- |
| Madagascar | *H. iboina* | 1.2918 | 0.0046 | 1.2803 | 1.3032 | A |
| Madagascar | *H. fraterna* | 0.8695 | 0.0064 | 0.8535 | 0.8854 | B |
| Africa | *B. martius* | 1.2456 | 0.0055 | 1.2319 | 1.2593 | a |
| Africa | *B. anynana* | 0.8013 | 0.0039 | 0.7917 | 0.8109 | b |

**Table S3:** (A) Analysis of deviance table for egg area corrected for body size. Body size was estimated as the average pupal weight of each of the species obtained from the common garden experiment. (B) model-estimated marginal means with upper and lower confidence limits. Significant differences (Tukey’s HSD, α = 0.05) between species were obtained using the R package *emmeans* and are indicated by different letters coding for each radiation independently.

**A.**
|  | Df | Deviance | P value |
| --- | --- | --- | --- |
| Habitat | 1 | 114.44 | <0.001 |
| Genus | 1 | 78.949 | <0.001 |
| Habitat*Genus | 1 | 12.151 | <0.001 |

**B.**
| Radiation | species | Mean | SE | LCL | UCL | Group |
| --- | --- | --- | --- | --- | --- | --- |
| Madagascar | *H. iboina* | 5.1568 | 0.0203 | 5.1060 | 5.2075 | A |
| Madagascar | *H. fraterna* | 4.9345 | 0.0283 | 4.8638 | 5.0051 | B |
| Africa | *B. martius* | 4.7578 | 0.0244 | 4.6970 | 4.8186 | a |
| Africa | *B. anynana* | 4.0328 | 0.0170 | 3.9902 | 4.0754 | b |

**Table S4:** (A) Analysis of deviance table for egg development time and (B) model-estimated marginal means with upper and lower confidence limits. Significant differences (Tukey’s HSD, α = 0.05) between species were obtained using the R package *emmeans* and are indicated by different letters coding for each radiation independently.

**A.**
|  | Df | Deviance | P value |
| --- | --- | --- | --- |
| Habitat | 1 | 256.331 | <0.001 |
| Genus | 1 | 22.381 | <0.001 |
| Habitat*Genus | 1 | 12.404 | <0.001 |

**B.**
| Radiation | Species | Mean | SE | LCL | UCL | Group |
| --- | --- | --- | --- | --- | --- | --- |
| Madagascar | *H. iboina* | 4.8301 | 0.0824 | 4.6292 | 5.0396 | A |
| Madagascar | *H. fraterna* | 3.9585 | 0.1084 | 3.6975 | 4.2378 | B |
| Africa | *B. martius* | 4.6597 | 0.1100 | 4.3936 | 4.9420 | a |
| Africa | *B. anynana* | 3.2713 | 0.0580 | 3.1300 | 3.4190 | b |

**Table S5:** (A) Analysis of deviance table for larval development time and (B) model-estimated marginal means with upper and lower confidence limits. Significant differences (Tukey’s HSD, α = 0.05) between species were obtained using the R package *emmeans* and are indicated by different letters coding for each radiation independently. (M=male, F=female).

**A.**
|  | Df | Deviance | P value |
| --- | --- | --- | --- |
| Habitat | 1 | 167.983 | <0.001 |
| Genus | 1 | 75.709 | <0.001 |
| Sex | 1 | 9.183 | 0.0024 |
| Habitat*Genus | 1 | 15.557 | <0.001 |
| Habitat*Sex | 1 | 0.275 | 0.5998 |
| Genus*Sex | 1 | 1.291 | 0.2557 |
| Habitat*Genus*Sex | 1 | 1.958 | 0.1617 |

**B.**
| Radiation | Species | Sex | Mean | SE | LCL | UCL | Group |
| --- | --- | --- | --- | --- | --- | --- | --- |
| Madagascar | *H. iboina* | M | 29.1320 | 0.7413 | 27.3426 | 31.0386 | A |
| Madagascar | *H. iboina* | F | 31.8214 | 0.6154 | 30.3246 | 33.3920 | B |
| Madagascar | *H. fraterna* | M | 25.6666 | 0.7817 | 23.7914 | 27.6896 | C |
| Madagascar | *H. fraterna* | F | 27.04 | 0.7353 | 25.2688 | 28.9352 | AC |
| Africa | *B. martius* | M | 37.0468 | 0.7608 | 35.1993 | 38.9913 | a |
| Africa | *B. martius* | F | 37.5428 | 0.7323 | 35.7622 | 39.4121 | a |
| Africa | *B. anynana* | M | 26.9803 | 0.7273 | 25.2281 | 28.8543 | b |
| Africa | *B. anynana* | F | 29.0491 | 0.6900 | 27.3801 | 30.8199 | b |

**Table S6:** (A) Analysis of deviance table for pupal development time and (B) model-estimated marginal means with upper and lower confidence limits. Significant differences (Tukey’s HSD, α = 0.05) between species were obtained using the R package *emmeans* and are indicated by different letters coding for each radiation independently. (M=male, F=female).

**A.**
|  | Df | Deviance | P value |
| --- | --- | --- | --- |
| Habitat | 1 | 88.684 | <0.001 |
| Genus | 1 | 0.653 | 0.4191 |
| Sex | 1 | 3.442 | 0.0635 |
| Habitat*Genus | 1 | 9.936 | 0.0016 |
| Habitat*Sex | 1 | 0.320 | 0.5715 |
| Genus*Sex | 1 | 0 | 0.9971 |
| Habitat*Genus*Sex | 1 | 0.007 | 0.9345 |

**B.**
| Radiation | Species | Sex | Mean | SE | LCL | UCL | Group |
| --- | --- | --- | --- | --- | --- | --- | --- |
| Madagascar | *H. iboina* | M | 9.9792 | 0.4560 | 8.9057 | 11.1821 | A |
| Madagascar | *H. iboina* | F | 9.3625 | 0.3421 | 8.5480 | 10.2546 | AB |
| Madagascar | *H. fraterna* | M | 8.0714 | 0.4384 | 7.0501 | 9.2407 | BC |
| Madagascar | *H. fraterna* | F | 7.8125 | 0.4034 | 6.8695 | 8.8849 | C |
| Africa | *B. martius* | M | 11.0000 | 0.4179 | 10.0069 | 12.0917 | A |
| Africa | *B. martius* | F | 10.2813 | 0.4008 | 9.3298 | 11.3297 | A |
| Africa | *B. anynana* | M | 7.2157 | 0.3761 | 6.3370 | 8.2162 | B |
| Africa | *B. anynana* | F | 7.0328 | 0.3395 | 6.2359 | 7.9315 | B |

**Table S7:** (A) Analysis of deviance table for total development time and (B) model-estimated marginal means with upper and lower confidence limits. Significant differences (Tukey’s HSD, α = 0.05) between species were obtained using the R package *emmeans* and are indicated by different letters coding for each radiation independently. (M=male, F=female).

**A.**
|  | Df | Deviance | P value |
| --- | --- | --- | --- |
| Habitat | 1 | 241.437 | <0.001 |
| Genus | 1 | 63.967 | <0.001 |
| Sex | 1 | 2.9885 | 0.0838 |
| Habitat*Genus | 1 | 25.14 | <0.001 |
| Habitat*Sex | 1 | 0.497 | 0.4807 |
| Genus*Sex | 1 | 0.911 | 0.3399 |
| Habitat*Genus*Sex | 1 | 1.841 | 0.1748 |

**B.**
| Radiation | Species | Sex | Mean | SE | LCL | UCL | Group |
| --- | --- | --- | --- | --- | --- | --- | --- |
| Madagascar | *H. iboina* | M | 38.7500 | 0.8985 | 36.5753 | 41.0540 | A |
| Madagascar | *H. iboina* | F | 41.0125 | 0.7160 | 39.2672 | 42.8353 | A |
| Madagascar | *H. fraterna* | M | 33.7381 | 0.8963 | 31.5778 | 36.0461 | B |
| Madagascar | *H. fraterna* | F | 34.8125 | 0.8516 | 32.7545 | 36.9998 | B |
| Africa | *B. martius* | M | 48.0159 | 0.8730 | 45.8898 | 50.2405 | a |
| Africa | *B. martius* | F | 47.7813 | 0.8640 | 45.6767 | 49.9827 | a |
| Africa | *B. anynana* | M | 34.1961 | 0.8188 | 32.2160 | 36.2978 | b |
| Africa | *B. anynana* | F | 36.0820 | 0.7691 | 34.2162 | 38.0495 | b |

**Table S8:** (A) Analysis of deviance table for pupal weight and (B) model-estimated marginal means with upper and lower confidence limits. Significant differences (Tukey’s HSD, α = 0.05) between species were obtained using the R package *emmeans* and are indicated by different letters coding for each radiation independently. (M=male, F=female).

**A.**
|  | Df | Deviance | P value |
| --- | --- | --- | --- |
| Habitat | 1 | 5526 | <0.001 |
| Genus | 1 | 3639 | <0.001 |
| Sex | 1 | 8611 | <0.001 |
| Habitat*Genus | 1 | 3926 | 0.0013 |
| Habitat*Sex | 1 | 1333 | <0.001 |
| Genus*Sex | 1 | 2155 | <0.001 |
| Habitat*Genus*Sex | 1 | 2300 | <0.001 |

**B.**
| Radiation | Species | Sex | Mean | SE | LCL | UCL | Group |
| --- | --- | --- | --- | --- | --- | --- | --- |
| Madagascar | *H. iboina* | M | 223.7358 | 2.6817 | 217.0559 | 230.4158 | A |
| Madagascar | *H. iboina* | F | 267.5310 | 2.1302 | 262.2249 | 272.8370 | B |
| Madagascar | *H. fraterna* | M | 153.5452 | 3.0125 | 146.0413 | 161.0491 | C |
| Madagascar | *H. fraterna* | F | 189.8800 | 2.7610 | 183.0026 | 196.7574 | D |
| Africa | *B. martius* | M | 266.2844 | 2.4404 | 260.2055 | 272.3632 | a |
| Africa | *B. martius* | F | 258.2657 | 2.3335 | 252.4532 | 264.0782 | b |
| Africa | *B. anynana* | M | 176.4000 | 2.7338 | 169.5903 | 183.2097 | c |
| Africa | *B. anynana* | F | 217.7689 | 2.4997 | 211.5423 | 223.9954 | d |

**Table S9:** (A) Analysis of deviance table for growth rates and (B) model-estimated marginal means with upper and lower confidence limits. Significant differences (Tukey’s HSD, α = 0.05) between species were obtained using the R package *emmeans* and are indicated by different letters coding for each radiation independently. (M=male, F=female).

**A.**
|  | Df | Deviance | P value |
| --- | --- | --- | --- |
| Habitat | 1 | 0.0855 | <0.001 |
| Genus | 1 | 0.0517 | <0.001 |
| Sex | 1 | 0.0030 | <0.001 |
| Habitat*Genus | 1 | 0.0175 | <0.001 |
| Habitat*Sex | 1 | 0.0001 | 0.5039 |
| Genus*Sex | 1 | 0.0002 | 0.3181 |
| Habitat*Genus*Sex | 1 | 0.0009 | 0.0482 |

**B.**
| Radiation | Species | sex | Mean | SE | LCL | UCL | Group |
| --- | --- | --- | --- | --- | --- | --- | --- |
| Madagascar | *H. iboina* | M | 0.1873 | 0.0021 | 0.1821 | 0.1926 | A |
| Madagascar | *H. iboina* | F | 0.1769 | 0.0017 | 0.1727 | 0.1810 | B |
| Madagascar | *H. fraterna* | M | 0.1980 | 0.0024 | 0.1921 | 0.2038 | C |
| Madagascar | *H. fraterna* | F | 0.1953 | 0.0022 | 0.1899 | 0.2007 | C |
| Africa | *B. martius* | M | 0.1511 | 0.0019 | 0.1464 | 0.1559 | a |
| Africa | *B. martius* | F | 0.1484 | 0.0018 | 0.1439 | 0.1530 | a |
| Africa | *B. anynana* | M | 0.1928 | 0.0021 | 0.1875 | 0.1981 | b |
| Africa | *B. anynana* | F | 0.1865 | 0.0020 | 0.1817 | 0.1914 | b |

**Table S10:** Generalised linear mixed models for testing variation in fecundity. (A) Minimum adequate model in bold and (B) analysis of deviance table for the minimum adequate model.

| Models | AIC |
| --- | --- |
| egg.no ~ pupal weight + habitat * genus * day + day<sup>2</sup> + (1\|female id) | 4356.22 |
| **egg.no ~ pupal weight+ habitat + genus + day + habitat\*genus + genus\*day + habitat\*day+ day<sup>2</sup>+ (1\|female id)** | **4354.25** |

**B.**
|  | Chisq | DF | P value |
| --- | --- | --- | --- |
| Pupal weight | 2.7387 | 1 | 0.0979 |
| Habitat | 20.1470 | 1 | <0.001 |
| Genus | 6.7569 | 1 | 0.0093 |
| Day | 47.2145 | 1 | <0.001 |
| Day<sup>2</sup> | 73.7138 | 1 | <0.001 |
| Habitat*Genus | 3.1830 | 1 | 0.0744 |
| Habitat*Day | 13.3650 | 1 | <0.001 |
| Genus*Day | 13.7207 | 1 | <0.001 |

**Figure S1:**
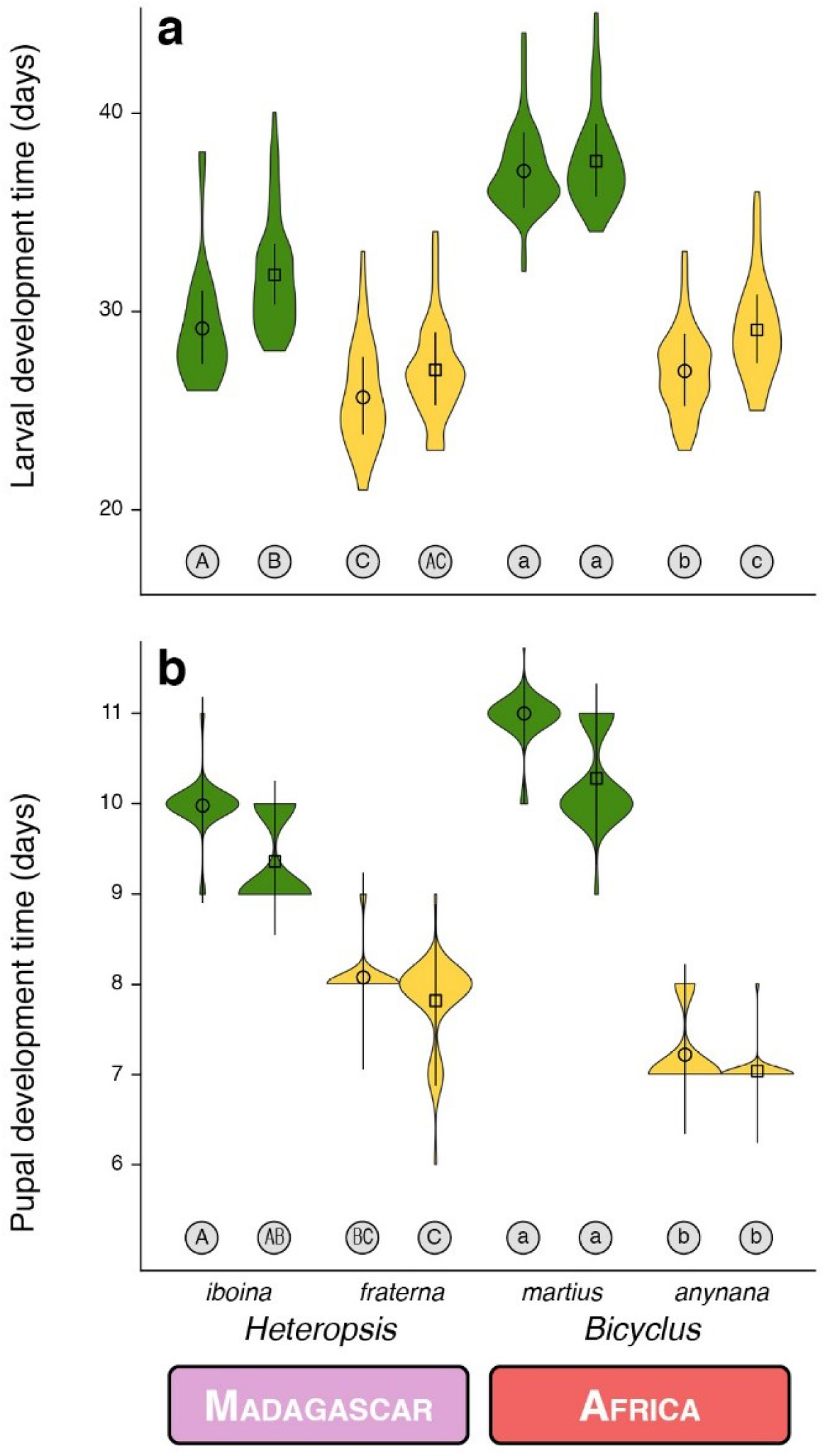
Violin plots with mean and 95% confidence intervals for the (a) larval and (b) pupal development time for replicate pairs of forest and savannah species from the Malagasy and African radiation reared in the common garden (green = forest species, yellow = savannah species). Sexes are denoted with different shapes (circles = males, squares = females) and significant differences in pairwise comparisons within each radiation are indicated by different letters coding for each radiation independently.

**Figure S2:**
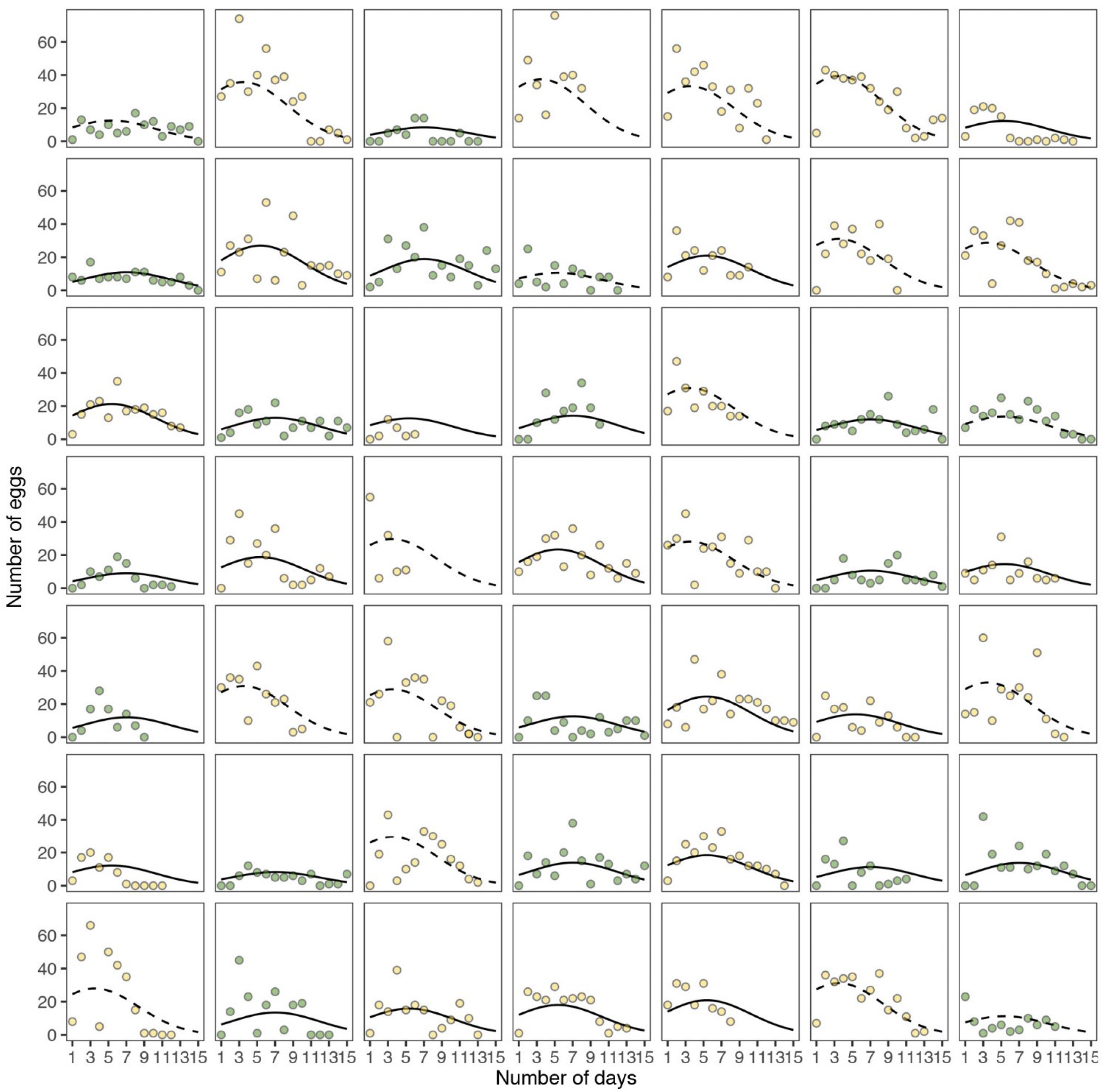
Fecundity curves for each female for replicate pairs of forest and savannah species from the Malagasy and African radiation (green = forest species, yellow = savannah species). Daily measurement of fecundity is shown in filled circles and the model estimated fecundity is shown as a line for each female. Dashed and solid lines represent species from the Malagasy and African radiation, respectively.

## Extracting data for species from the Asian radiation from published studies

We extracted data for three Australian species representing the Asian radiation from published studies (Braby & Jones 1994; Braby & Jones 1995). When possible, we extracted values from the plots showing mean and error bars as standard error using the WebPlotDigitiser Ver. 4.2. (https://automeris.io/WebPlotDigitizer/) (Rohtagi 2019), or used values provided in the tables. For some traits such as the total development time, information on confidence interval was not available.

**Figure S3:**
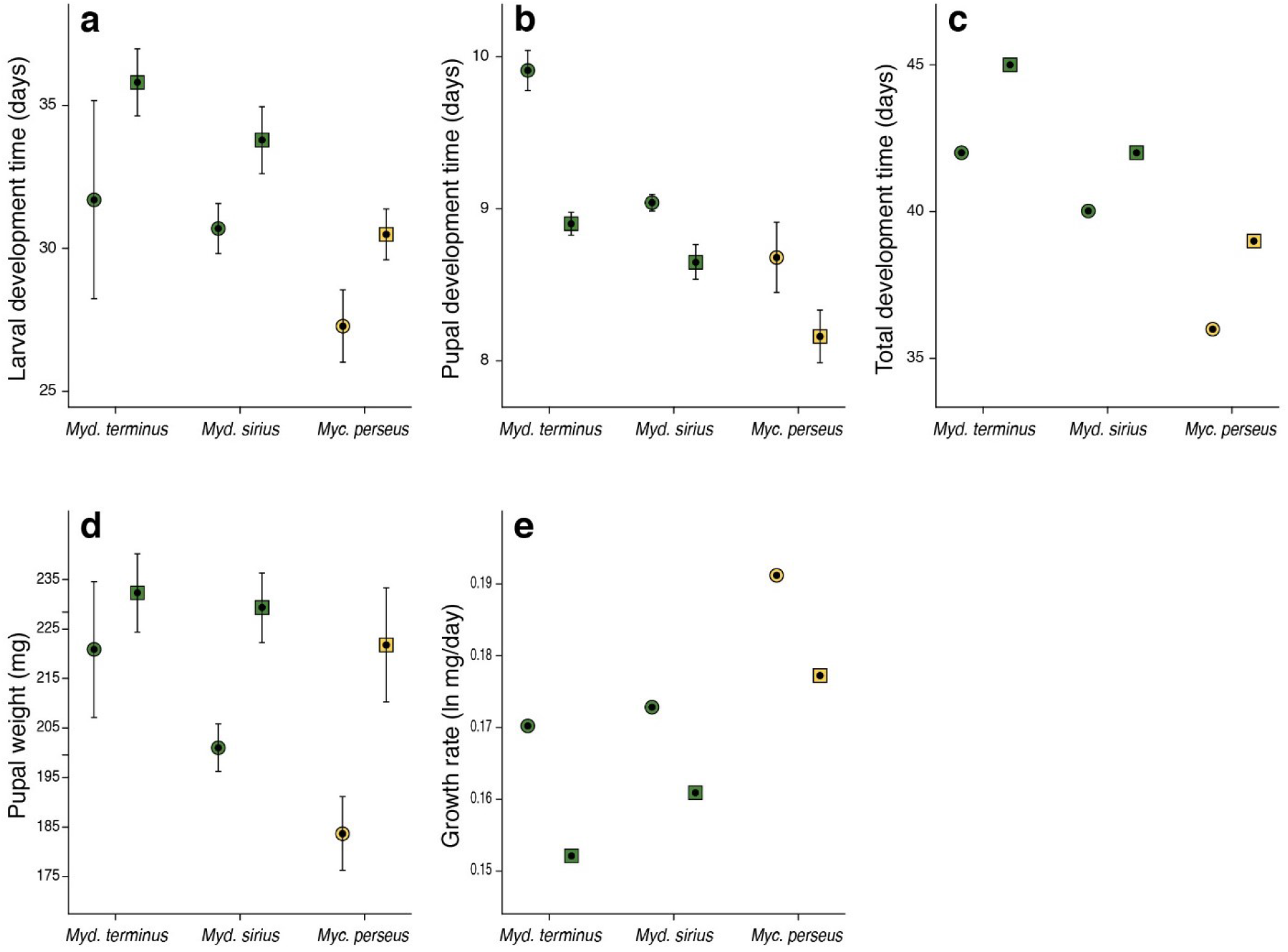
Mean values with 95% confidence intervals (a) larval and (b) pupal development time and (d) pupal weight for three species from the Asian radiation (green = forest species, yellow = savannah species). For the (c) total development time, only the mean value was available. (e) Growth rate was calculated as ln(pupal weight)/larval development time. Data was extracted from Braby & Jones (1994). Sexes are denoted with different shapes (circles = males, squares = females).

**Figure S4:**
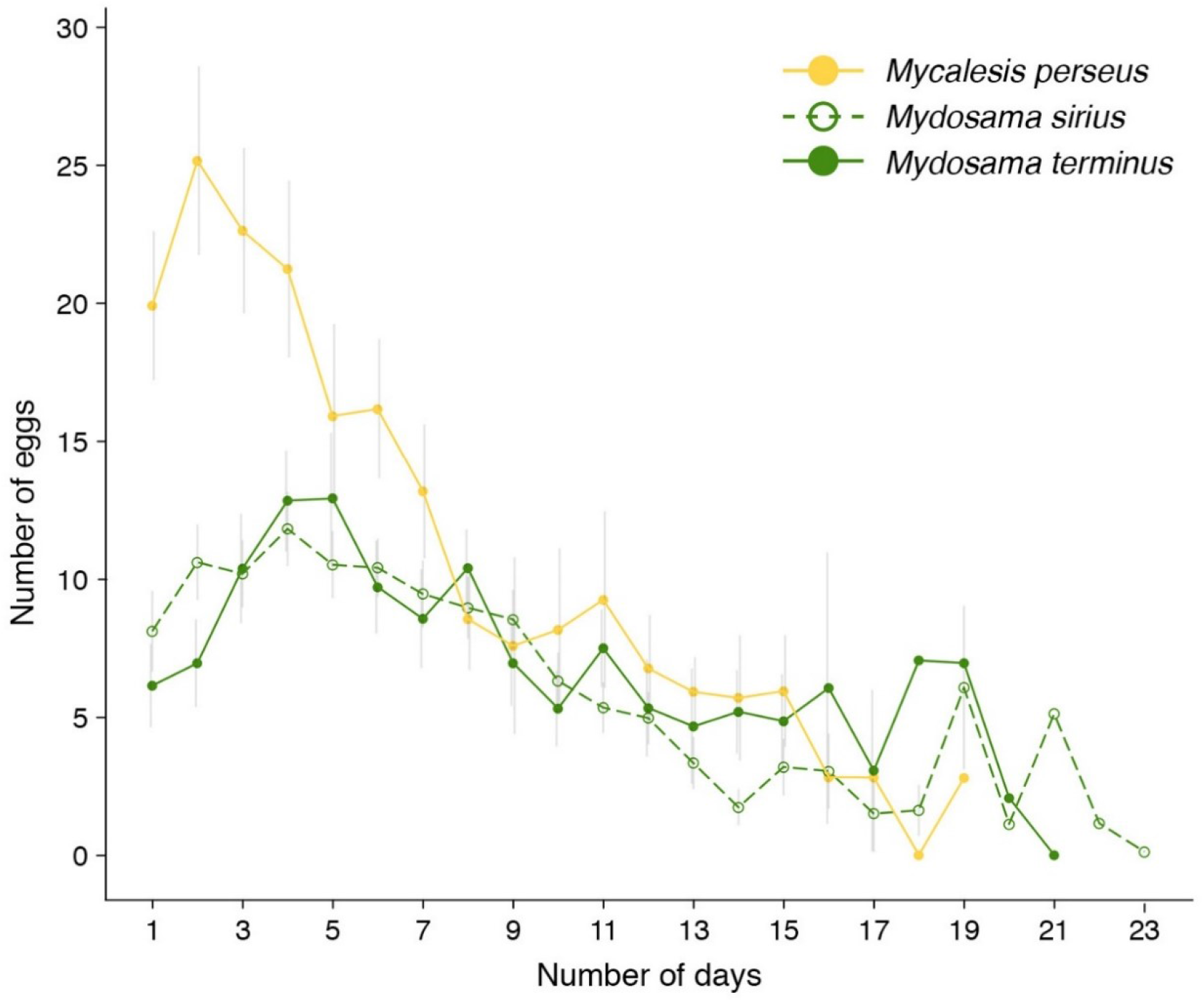
Fecundity curves (mean number of eggs ± SE) for three species from the Asian radiation (green = forest species, yellow = savannah species). Data was extracted from Braby & Jones (1995).

